# Two-dimensional video-based analysis of human gait using pose estimation

**DOI:** 10.1101/2020.07.24.218776

**Authors:** Jan Stenum, Cristina Rossi, Ryan T. Roemmich

**Author notes:** corresponding author: Contact information: G-04 Kennedy Krieger Institute, 707 North Broadway, Baltimore, MD 21205. Twitter: Jan Stenum – @janstenum, Cristina Rossi – @cristina_rossi, Ryan Roemmich – @RyanRoemmich.

## Abstract

Walking is the primary mode of human locomotion. Accordingly, people have been interested in studying human gait since at least the fourth century BC. Human gait analysis is now common in many fields of clinical and basic research, but gold standard approaches – e.g., three-dimensional motion capture, instrumented mats or footwear, and wearables – are often expensive, immobile, data-limited, and/or require specialized equipment or expertise for operation. Recent advances in video-based pose estimation have suggested exciting potential for analyzing human gait using only two-dimensional video inputs collected from readily accessible devices (e.g., smartphones, tablets). However, we currently lack: 1) data about the accuracy of video-based pose estimation approaches for human gait analysis relative to gold standard measurement techniques and 2) an available workflow for performing human gait analysis via video-based pose estimation. In this study, we compared a large set of spatiotemporal and sagittal kinematic gait parameters as measured by OpenPose (a freely available algorithm for video-based human pose estimation) and three-dimensional motion capture from trials where healthy adults walked overground. We found that OpenPose performed well in estimating many gait parameters (e.g., step time, step length, sagittal hip and knee angles) while some (e.g., double support time, sagittal ankle angles) were less accurate. We observed that mean values for individual participants – as are often of primary interest in clinical settings – were more accurate than individual step-by-step measurements. We also provide a workflow for users to perform their own gait analyses and offer suggestions and considerations for future approaches.

## INTRODUCTION

Humans have been interested in studying the walking patterns of animals and other humans for centuries, dating back to Aristotle in the fourth century BC (see Baker (2007) for a detailed history of gait analysis). Gait analysis technologies have evolved from Borelli’s use of staggered poles to study his own gait to modern tools that include three-dimensional motion capture, instrumented gait mats, and a variety of wearable devices. Although technological advances continue to expand our abilities to measure human walking in clinical and laboratory settings, many limitations persist. Current techniques remain expensive, are often time consuming, and require specialized equipment or expertise that is often not widely accessible.

Recent progress in video-based pose estimation has enabled automated analysis of the movements of human (Andriluka et al., 2014; Toshev and Szegedy, 2014; Insafutdinov et al., 2016; Pishchulin et al., 2016; Martinez et al., 2017; Cao et al., 2019) and animals (Mathis et al., 2018; Nath et al., 2019) using only digital video inputs. The learning algorithms at the core of human pose estimation approaches use networks that are generally trained on many images of different people (e.g., MPII (Andriluka et al., 2014) and COCO (Lin et al., 2014) datasets), resulting in robust networks capable of detecting keypoints (e.g., body landmarks) in new images beyond the training dataset. These software packages are freely available and have the potential to expand the ability to generate large datasets of human gait data by enabling data collection in any setting (including the home or clinic) with little cost of time, money, or effort.

In our view, three central barriers have precluded wide use of pose estimation for human gait analysis. First, interest in pose estimation is relatively new and has been primarily confined to developers and researchers within the computer science, neuroscience, and engineering communities. Second, there is a need for validation of gait parameters as calculated by pose estimation approaches against gold standard methods for measuring human gait data (e.g., three-dimensional motion capture systems). Third, there is a need for dissemination of a workflow that produces spatiotemporal and kinematic gait parameter outputs from a simple digital video input.

The goals of this study were two-fold: 1) compare spatiotemporal and kinematic gait parameters as measured by three-dimensional motion capture and pose estimation via OpenPose, and 2) provide a workflow and suggestions for performing automated human gait analysis from digital video. OpenPose is a freely available human pose estimation algorithm that uses Part Affinity Fields to detect up to 135 keypoints in images of humans (Martinez et al., 2017; Cao et al., 2019). Several prior studies have used OpenPose to study certain features of human walking (Chambers et al., 2019; Sato et al., 2019; Viswakumar et al., 2019; Zago et al., 2020), but there remain important needs for robust validation of a comprehensive set of gait parameters against motion capture measurements and a shareable workflow. First, we used OpenPose to detect keypoints on videos of healthy adults walking overground. These videos were provided in a freely available dataset that includes synchronized digital videos and three-dimensional motion capture gait data (Kwolek et al., 2019). We then developed a workflow to calculate a wide variety of spatiotemporal and kinematic gait parameters from the OpenPose outputs. We validated these parameters against measurements calculated from the motion capture data and across different camera views within the same walking trials to test the robustness of using OpenPose to estimate gait parameters from different camera viewpoints.

We wish to state upfront that the workflow provided here is simply one approach to human gait analysis using pose estimation. We considered it important to test OpenPose in an “out of the box” approach by using the available demo without modification, as we anticipate that this is how the software is most likely to be used by potential users who may have interest in performing gait analysis but have relatively little expertise in computer science or engineering. While our results show that this workflow provides, in our view, reasonably accurate estimates of many gait parameters, there is almost certainly opportunity to improve upon these results by adjusting OpenPose parameters, using other pose estimation algorithms, or using other methods of video recording. We do not claim to provide an optimized approach to human gait analysis through video-based pose estimation, but rather an approach that we found to be easy to use, fast, and reasonably accurate.

## METHODS

### Participants

We used a publicly available dataset (Kwolek et al., 2019) of overground walking sequences from 32 healthy participants (10 men and 22 women) made available at http://bytom.pja.edu.pl/projekty/hm-gpjatk/. The dataset included synchronized three-dimensional motion capture files and digital video recordings of the walking sequences. The dataset does not contain identifiable participant information and faces have been blurred in the video recordings. Our analyses of these previously published videos were deemed exempt by the Johns Hopkins Institutional Review Board.

### Walking sequences

The laboratory space in which the participants walked included ten motion capture cameras and four video cameras that recorded left and right side sagittal plane views and front and back frontal plane views (see Figure 1A for overview of laboratory space). We used a subset of the data (sequences labelled s1) that consisted of a single walking bout of about five meters that included gait initiation and termination. We excluded data for one participant because the data belonged to another subset with diagonal walking sequences. Therefore, we analyzed 31 total gait trials (one trial per participant). The mean±SD duration of the video recordings was 5.12±0.73 seconds.

**Figure 1.**
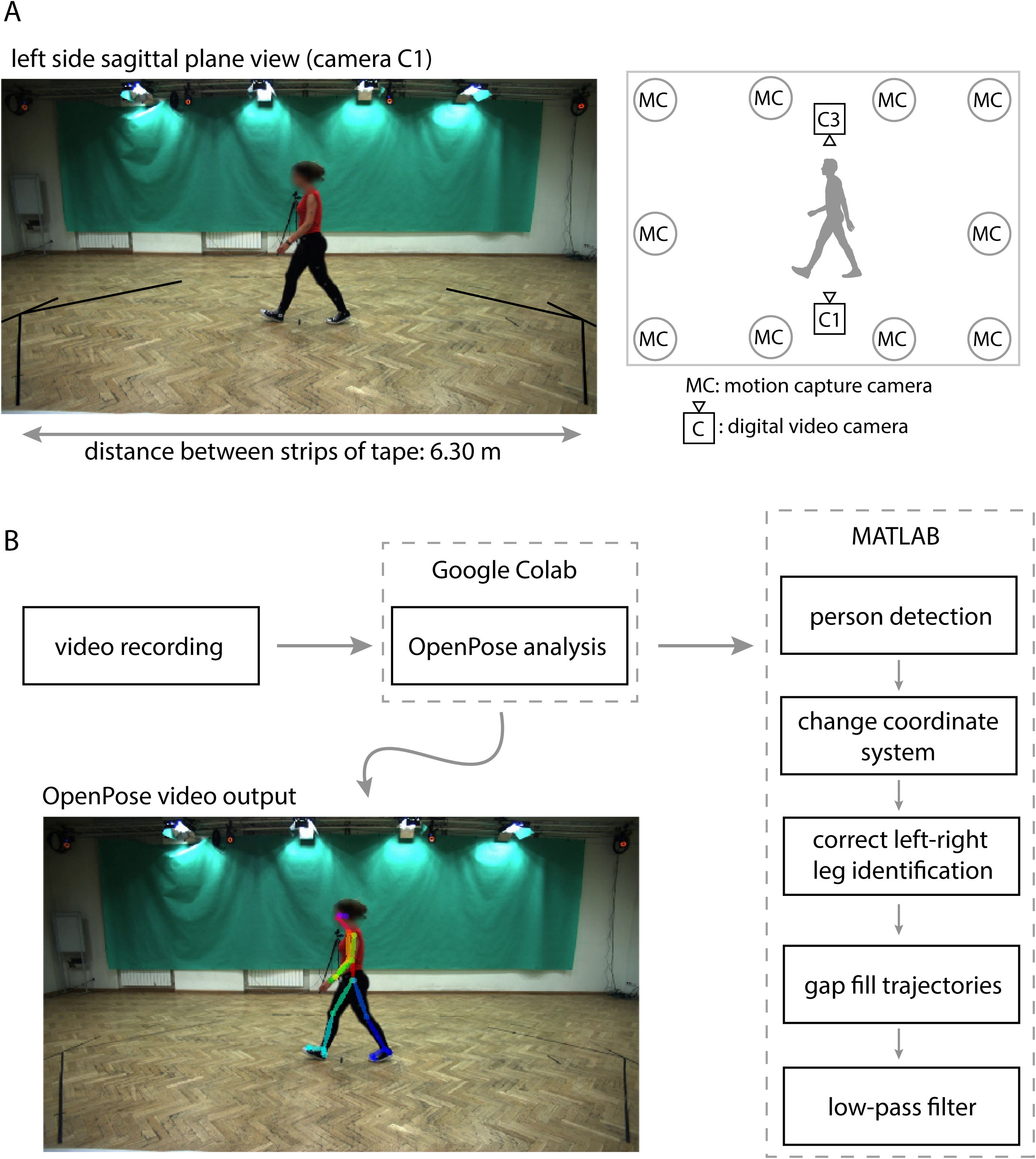
Overview of laboratory space and workflow. A) Representative image frame of original video recording from left side sagittal plane view with diagram of motion capture and video cameras. We used a known distance of 6.30 m between two strips of tape on the floor to dimensionalize pixel coordinates of OpenPose keypoints. The public dataset (Kwolek et al, 2019) that we used is made available at http://bytom.pja.edu.pl/projekty/hm-gpjatk/. See Release Agreement for copyrights and permissions. B) Workflow of video recordings (available at https://github.com/janstenum/GaitAnalysis-PoseEstimation). We analyzed video recording with OpenPose using Google Colab and next processed the data using custom MATLAB scripts.

### Data collection

The motion capture cameras (Vicon, MX-T40) recorded three-dimensional marker positions at 100 Hz. Motion was recorded by tracking markers that were placed on the seventh cervical vertebrae (C7), tenth thoracic vertebrae (T10), manubrium, sternum, right upper back and bilaterally on the front and back of the head, shoulder, upper arm, elbow, forearm, wrist (at radius and ulna), middle finger, anterior superior iliac spine (ASIS), posterior superior iliac spine (PSIS), thigh, knee, shank, ankle, heel and toe.

Four video cameras recorded left (camera C1) and right side (camera C3) sagittal plane views and front (camera C4) and back (camera C2) frontal plane views of the walking sequences at 25 Hz. We only analyzed video recordings of left and right side sagittal plane views (cameras C1 and C3). The digital camera images were RBG files with 960×540 pixel resolution. Motion capture and video recording were synchronized so that the time of every fourth motion capture data point corresponded to each time point of the video frames. Cameras were mounted on tripods and the height was set about 1.3 m. The distance from cameras C1 or C3 to the participants was about 3.2 m.

### Data processing

Motion capture data had already been smoothed and were therefore not processed further. We used the following workflow to process sagittal plane video recordings and obtain two-dimensional coordinates of OpenPose keypoints. We first analyzed the video recordings with OpenPose using our provided Google Colaboratory notebook and next processed the data using custom written MATLAB software (also provided). The workflow is shown in Figure 1B and is as follows (detailed instructions, sample videos, and software for all steps can be found at https://github.com/janstenum/GaitAnalysis-PoseEstimation):

1. OpenPose analyses
  a. ii) We used Google Colaboratory to run OpenPose analyses of the sagittal plane video recordings. Google Colaboratory executes Python code and allows the user to access GPUs remotely through Google cloud services. This allows for much faster analysis than can be executed on a CPU. Note that it was possible to use Google Colaboratory for our study because we analyzed publicly available videos; users that aim to analyze videos of research participants or patients may need to run the software locally through a Python environment to avoid uploading identifiable participant or patient information into Google Drive if this is not deemed sufficiently secure by the user’s institution. In our Google Colaboratory notebook, we also provide code for analyzing YouTube videos with OpenPose should this be of interest to some users.
  b. iii) Video recordings were analyzed in OpenPose using the BODY_25 keypoint model that tracks the following 25 keypoints: nose, neck, midhip and bilateral keypoints at eyes, ears, shoulders, elbows, wrists, hips, knees, ankles, heels, big toes and small toes.
  c. iv) The outputs of the OpenPose analyses yielded: 1) JSON files for every video frame containing pixel coordinates (origin at upper left corner of the video) of each keypoint detected in the frame, and 2) a new video file in which a stick figure that represents the detected keypoints is overlaid onto the original video recording. The JSON files were then downloaded for further offline analysis in MATLAB.
2. MATLAB processing steps
  a. In about 20% of the total number of frames (3,970), OpenPose detected false positive persons (e.g., from an anatomy poster that was visible in some videos or from tripods that were visible). We visually inspected all frames in which multiple persons were detected so that the keypoints tracked were always the keypoints from the participant. OpenPose consistently assigned the participant as person ID #1 in all frames wherein multiple persons were detected. We have since updated our Google Colab notebook to limit OpenPose to identification of one person per video. As a result, we have not provided this step in our workflow.
  b. We changed the pixel coordinate system so that positive vertical was directed upward and positive horizontal was in the direction of travel. The location of the origin depended on left (camera C1) or right side (camera C3) views of the video recording. For the left side view, the origin was set at the lower right corner; for the right side view, the origin was set at the lower left corner. Note that all our further analyses were invariant to the location of the origin.
  c. In approximately 5% of the total frames, OpenPose erroneously switched the left-right identification of the limbs. We visually inspected antererior-posterior trajectories of the left and right ankle to identify and correct (hip, knee, ankle, heel, big toe and small toe keypoints) these frames. We also corrected frames in which keypoints on the left and right legs were identified on the same leg.
  d. We filled gaps in keypoint trajectories (i.e., frames where OpenPose did not detect all keypoints) using linear interpolation for gaps spanning up to 120 ms (i.e., for gaps spanning up to two video frames).
  e. We filtered trajectories using zero-lag 4^th^ order low-pass Butterworth filter with a cut-off frequency at 5 Hz.
  f. Last, we calculated a scaling factor to obtain dimensionalized coordinate values from the pixel coordinates. The scaling factor (*s*) was calculated as:

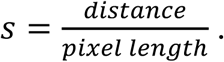

We used the distance between two strips of tapes on the floor at each end of the walkway that ran parallel to the viewpoints of cameras C1 and C3 (see Figure 1A). Pixel length was taken as the horizontal pixel length between the midpoints of each strip of tape. We then calculated dimensionalized coordinates as:

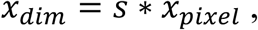

And

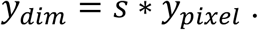

The distance between the strips of tape was not measured during the original data collection for the dataset and the tape has since been removed from the floor. We therefore estimated the distance between the strips of tape by: 1) calculating individual scaling factors from each participant based on the horizontal distance traversed by left and right ankle markers in motion capture data and left and right ankle keypoints in OpenPose data; 2) calculating the distance between the strips of tape for each individual participant; 3) calculating the ensemble mean distance. The ensemble mean distance was 6.30 m and we used this fixed value to calculate scaling factors for each individual participant. To examine how robustly the known distance was estimated, we also calculated the distance between the strips of tape by using the ‘CLAV’ marker (placed on the manubrium) from the motion capture dataset and the neck marker in OpenPose and obtained an ensemble mean distance of 5.91 m (2.5% reduction relative to the distance of 6.30 m that we used here). This means that there is a margin of uncertainty associated with the scaling factor that we used to dimensionalize the pixel coordinates obtained with OpenPose. Note that the use of the scaling factor assumes that the participants walked perpendicularly and at a fixed depth relative to the cameras. Barring natural side-to-side fluctuations in gait, we find that this is a reasonable assumption since net displacement of medio-lateral position of the C7 marker in the motion capture data from the start to the end of the walking sequence was (ensemble mean±SD) –0.025±0.081 m.

We calculated event timings of left and right heel-strikes and toe-offs in motion capture data and data of OpenPose left (C1) and right (C3) side views by independently applying the same method to each set of data (Zeni et al., 2008). Heel-strikes and toe-offs were defined by the time points of positive and negative peaks of the anterior-posterior ankle trajectories relative to the pelvis (midpoint of left and right ASIS and PSIS markers in motion capture data; midhip keypoint in OpenPose data).

We calculated the following spatiotemporal gait parameters in the motion capture and OpenPose data:

- Step time: duration in seconds between consecutive bilateral heel-strikes.
- Stance time: duration in seconds between heel-strike and toe-off of the same leg.
- Swing time: duration in seconds between toe-off and heel-strike of the same leg.
- Double support time: duration in seconds between heel-strike of one leg and toe-off of the contralateral leg.
- Step length: anterior-posterior distance in meters between left and right ankle markers (motion capture) or ankle keypoints (OpenPose) at heel-strike.
- Gait speed: step length divided by step time.

For step time and double support time, right step refers to the duration until right heel-strike and vice versa for the left step. For step length, right step refers to the distance between the ankles at right heel-strike and vice versa for the left step. We calculated step time, stance time, swing time, double support time and step length for all steps and as averages for individual participants. Gait speed was calculated from individual participant means of step time and step length.

We calculated sagittal plane hip, knee and ankle angles of left and right legs using two-dimensional marker (motion capture) and keypoint (OpenPose) coordinates. The motion capture dataset did not contain hip markers, so we created virtual hip markers in the motion capture data by using the midpoints of the ASIS and PSIS markers from each leg. We used the following markers or keypoints to calculate joint angles: virtual hip (motion capture) or hip (OpenPose) and knee (hip angle); hip, knee and ankle (knee angle); knee, ankle, toe (motion capture) or big toe (OpenPose) (ankle angle). All joint angles were calculated as the mean across the stride cycle of individual participants. We calculated cross-correlations at time lag zero to assess the similarity between mean joint angle trajectories (i.e., one measure per participant) calculated by motion capture and OpenPose left (C1) side view, by motion capture and OpenPose right (C3) side view, and by OpenPose left (C1) and right (C3) side views. Last, we calculated peak hip flexion and extension, peak knee flexion and extension, and peak ankle dorsiflexion and plantarflexion.

### Statistical analyses

We obtained gait event (i.e., heel-strike and toe-off) times, spatiotemporal gait parameters and sagittal joint angles from three measurement systems: motion capture, OpenPose left (C1) and right (C3) side views. We used one-way repeated measures ANOVA to assess potential differences in gait event times, gait parameters, and peak joint angles among measurement systems. In the event of a statistically significant main effect, we performed post-hoc pairwise comparisons with Bonferroni corrections. We calculated Pearson correlation coefficients (*r*) and intra-class correlation coefficients (ICC_C-1_ and ICC_A-1_) of spatiotemporal gait parameters and peak joint angles to assess correlations (*r*), consistency (ICC_C-1_) and agreement (ICC_A-1_) between 1) motion capture and OpenPose left (C1) side view, 2) motion capture and OpenPose right (C3) side view and 3) OpenPose left (C1) and right (C3) side views. We set the level of significance at 0.05 for all analyses.

## RESULTS

### Event times

First, we examined how well OpenPose identified common gait events (i.e., heel-strikes and toe-offs). Differences in event times as identified by motion capture, OpenPose left (C1) views, and OpenPose right (C3) views are shown in Table 1. We did not observe significant differences in event times as calculated by motion capture and OpenPose with the exception of motion capture vs. OpenPose right (C3) view in estimating left and right toe-off (post hoc *P*≤0.040). Mean differences between motion capture and OpenPose left (C1) and right (C3) side views were less than 0.010 s for heel-strikes and less than 0.020 s for toe-offs – durations less than one and two motion capture frames, respectively. For all variables reported in this study, standard deviations and ranges of discrepancies between the measurement systems are reported in their respective tables to provide insight into the distributions of the discrepancies.

**Table 1.** Differences in event times for all steps. *MC*: motion capture; *C1*: OpenPose left side view; *C3*: OpenPose right side view. Asterisks (*) denote *P* < 0.05.

|  | N | mean±SD |  |  | range |  |  | F | post-hoc (P) |  |  |
| --- | --- | --- | --- | --- | --- | --- | --- | --- | --- | --- | --- |
|  |  | MC-C1 | MC-C3 | C1-C3 | MC-C1 | MC-C3 | C1-C3 |  | MC-C1 | MC-C3 | C1-C3 |
| Left Heel-Strike Time (s) | 107 | -0.006±0.020 | -0.005±0.018 | 0.001±0.023 | [-0.050, 0.060] | [-0.050, 0.040] | [-0.080, 0.040] | 3.5* | 1.000 | 0.227 | 0.013 |
| Right Heel-Strike Time (s) | 109 | -0.005±0.016 | -0.006±0.019 | -0.001±0.022 | [-0.040, 0.040] | [-0.050, 0.030] | [-0.040, 0.040] | 2.7 | - | - | - |
| Left Toe-Off Time (s) | 109 | -0.014±0.029 | -0.014±0.025 | 0.000±0.035 | [-0.090, 0.060] | [-0.080, 0.040] | [-0.120, 0.080] | 8.0* | 1.000 | 0.040 | <0.001 |
| Right Toe-Off Time (s) | 107 | -0.016±0.024 | -0.011±0.027 | 0.005±0.030 | [-0.110, 0.050] | [-0.070, 0.070] | [-0.040, 0.120] | 21.8* | 0.226 | <0.001 | <0.001 |

When comparing OpenPose estimates from different viewpoints, we observed statistically significant differences in left heel-strike, left toe-off, and right toe-off times between OpenPose left (C1) and right (C3) side views (post-hoc *P*≤0.013). However, the magnitudes of the differences in event times between OpenPose left (C1) and right (C3) side views were generally small: less than 0.010 s on average.

### Spatiotemporal gait parameters

We next examined how well OpenPose estimated the following spatiotemporal gait parameters: step time, stance time, swing time, double support time, step length, and gait speed. In our first analysis, we investigated these parameters on an individual step basis (i.e., we included all steps across all participants in the analysis). In our second analysis, we investigated the same parameters on an individual mean basis (i.e., one mean value per participant).

#### Temporal parameters – all steps

When comparing temporal parameters between motion capture and OpenPose, we observed a significant main effect of measurement system on stance time, swing time, and double support time (post-hoc *P*≤0.001), but not step time, when analyzing all individual steps (Figure 2, Tables 2 and 3). We observed that OpenPose tended to overestimate stance time and double support time but underestimate swing time when compared to motion capture (Table 2). The magnitudes of these differences were again typically small, as mean differences were not larger than 0.010 s (i.e., one motion capture frame) for any parameter. Pearson and intra-class correlation coefficients between motion capture and OpenPose (C1 and C3) were strong for step time (Figure 2A), stance time (Figure 2B) and swing time (Figure 2C; all ≥0.839) but somewhat less strong for double support time (Figure 2D; all ≥0.660). All correlations were statistically significant (Table 3).

**Table 2.** Gait parameters calculated for all steps. *MC*: motion capture; *C1*: OpenPose left side view; *C3*: OpenPose right side view.

|  | N | mean±SD |  |  | mean±SD |  |  | range |  |  |
| --- | --- | --- | --- | --- | --- | --- | --- | --- | --- | --- |
|  |  | MC | C1 | C3 | MC-C1 | MC-C3 | C1-C3 | MC-C1 | MC-C3 | C1-C3 |
| Step Time (s) | 185 | 0.608±0.076 | 0.608±0.081 | 0.607±0.078 | -0.001±0.023 | 0.000±0.021 | 0.001±0.033 | [-0.060, 0.080] | [-0.060, 0.060] | [-0.120, 0.080] |
| Stance Time (s) | 154 | 0.746±0.099 | 0.755±0.105 | 0.755±0.102 | -0.009±0.027 | -0.009±0.024 | 0.000±0.033 | [-0.100, 0.060] | [-0.060, 0.050] | [-0.080, 0.120] |
| Swing Time (s) | 154 | 0.457±0.045 | 0.447±0.050 | 0.448±0.047 | 0.010±0.026 | 0.009±0.025 | -0.001±0.030 | [-0.060, 0.070] | [-0.050, 0.070] | [-0.080, 0.080] |
| Double Support Time (s) | 154 | 0.139±0.030 | 0.147±0.036 | 0.147±0.038 | -0.008±0.026 | -0.009±0.026 | -0.001±0.036 | [-0.080, 0.050] | [-0.070, 0.070] | [-0.080, 0.080] |
| Step Length (m) | 215 | 0.598±0.095 | 0.606±0.118 | 0.598±0.114 | -0.008±0.059 | 0.000±0.059 | 0.008±0.094 | [-0.106, 0.202] | [-0.112, 0.204] | [-0.282, 0.253] |

**Table 3.** Gait parameter calculated for all steps. *MC*: motion capture; *C1*: OpenPose left side view; *C3*: OpenPose right side view. Asterisks (*) denote *P* < 0.05.

|  | *F* | post-hoc (*P*) |  |  | *r* |  |  | ICC<sub>C-1</sub> |  |  | ICC<sub>A-1</sub> |  |  |
| --- | --- | --- | --- | --- | --- | --- | --- | --- | --- | --- | --- | --- | --- |
|  |  | *MC v C1* | *MC v C3* | *C1 v C3* | *MC v C1* | *MC v C3* | *C1 v C3* | *MC v C1* | *MC v C3* | *C1 v C3* | *MC v C1* | *MC v C3* | *C1 v C3* |
| Step Time | 0.2 | - | - | - | 0.957* | 0.963* | 0.911* | 0.955* | 0.962* | 0.911* | 0.955* | 0.963* | 0.911* |
| Stance Time | 9.7* | <0.001 | <0.001 | 1.000 | 0.967* | 0.971* | 0.951* | 0.966* | 0.971* | 0.950* | 0.962* | 0.967* | 0.950* |
| Swing Time | 12.3* | <0.001 | <0.001 | 1.000 | 0.860* | 0.858* | 0.810* | 0.855* | 0.857* | 0.809* | 0.839* | 0.842* | 0.810* |
| Double Support Time | 8.0* | 0.001 | <0.001 | 1.000 | 0.691* | 0.735* | 0.523* | 0.678* | 0.714* | 0.522* | 0.660* | 0.694* | 0.524* |
| Step Length | 1.6 | - | - | - | 0.869* | 0.857* | 0.671* | 0.849* | 0.843* | 0.671* | 0.848* | 0.844* | 0.671* |

**Figure 2.**
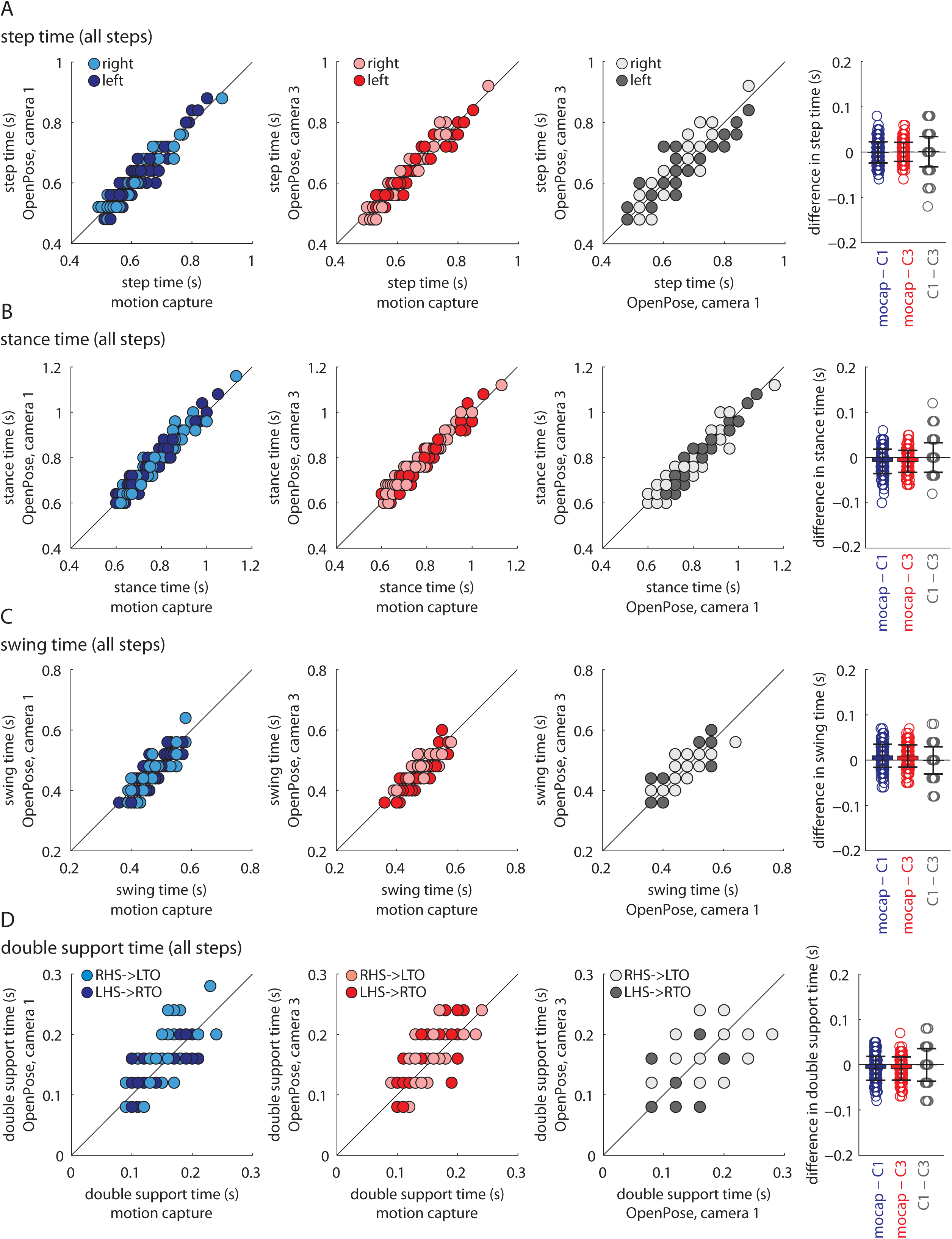
Temporal gait parameters for all individual steps shown for all participants and measurement systems: A) step time, B) stance time, C) swing time, and D) double support time. Data shown in blue represent comparisons between motion capture and OpenPose left side (C1) views, data shown in red represent comparisons between motion capture and OpenPose right side (C3) views, and data shown in gray represent comparisons between the two OpenPose views (C1 and C3). Dark circles represent left leg data, light circles represent right leg data. Color schemes and shading are consistent across all similar figures (Figures 3, 4, 8, 9, and 10). Bar plots on the far right show individual data, group means, and standard deviations to visualize the distribution of the differences observed between the measurement systems. Results from statistical analyses are shown in Table 3.

When we compared temporal gait parameters for all steps estimated with OpenPose (C1 and C3) from left (C1) and right (C3) side views, we did not observe statistically significant differences in any parameters (Table 3). Pearson and intra-class correlation coefficients were again strong for step time (Figure 2A), stance time (Figure 2B) and swing time (Figure 2C; all ≥0.809) but less strong for double support time (Figure 2D; all between 0.522 and 0.524). All correlations were statistically significant (Table 3).

#### Temporal parameters – individual participant means

When we compared temporal parameters between motion capture and OpenPose (C1 and C3) using individual participant means (as are often of interest clinically), we observed similar results except correlations between measurement systems became stronger for all parameters.

We again observed a significant main effect of measurement system on stance time, swing time, and double support time (post-hoc *P*≤0.003), but not step time (Figure 3, Tables 4 and 5). As expected based on our prior findings, we observed that OpenPose tended to overestimate stance time and double support time but underestimate swing time (Table 4). We observed strong Pearson and intra-class correlation coefficients for all temporal parameters (all ≥0.893). All correlations were statistically significant (Table 5).

**Table 4.** Gait parameters calculated as individual participant means. *MC*: motion capture; *C1*: OpenPose left side view; *C3*: OpenPose right side view.

|  | N | mean±SD |  |  | mean±SD |  |  | range |  |  |
| --- | --- | --- | --- | --- | --- | --- | --- | --- | --- | --- |
|  |  | MC | C1 | C3 | MC-C1 | MC-C3 | C1-C3 | MC-C1 | MC-C3 | C1-C3 |
| Step Time (s) | 31 | 0.607±0.068 | 0.608±0.068 | 0.607±0.067 | -0.001±0.004 | 0.001±0.005 | 0.001±0.007 | [-0.010, 0.010] | [-0.012, 0.010] | [-0.017, 0.013] |
| Stance Time (s) | 31 | 0.745±0.092 | 0.753±0.092 | 0.753±0.092 | -0.009±0.012 | -0.008±0.012 | 0.000±0.013 | [-0.036, 0.010] | [-0.030, 0.008] | [-0.024, 0.024] |
| Swing Time (s) | 31 | 0.457±0.043 | 0.447±0.043 | 0.448±0.040 | 0.010±0.012 | 0.009±0.013 | -0.001±0.012 | [-0.010, 0.034] | [-0.015, 0.033] | [-0.020, 0.024] |
| Double Support Time (s) | 31 | 0.138±0.028 | 0.146±0.027 | 0.146±0.028 | -0.008±0.012 | -0.008±0.013 | 0.000±0.013 | [-0.034, 0.014] | [-0.033, 0.015] | [-0.032, 0.020] |
| Step Length (m) | 31 | 0.603±0.060 | 0.611±0.063 | 0.603±0.065 | -0.008±0.020 | 0.000±0.015 | 0.008±0.023 | [-0.050, 0.031] | [-0.030, 0.038] | [-0.052, 0.057] |
| Gait Speed (m s<sup>-1</sup>) | 31 | 1.00±0.15 | 1.02±0.16 | 1.01±0.16 | -0.01±0.04 | 0.00±0.03 | 0.01±0.04 | [-0.09, 0.05] | [-0.06, 0.07] | [-0.09, 0.08] |

**Table 5.** Gait parameters calculated as individual participant means. *MC*: motion capture; *C1*: OpenPose left side view; *C3*: OpenPose right side view. Asterisks (*) denote *P* < 0.05.

|  | *F* | *post-hoc (P)* |  |  | *r* |  |  | *ICC<sub>C-1</sub>* |  |  | *ICC<sub>A-1</sub>* |  |  |
| --- | --- | --- | --- | --- | --- | --- | --- | --- | --- | --- | --- | --- | --- |
|  |  | *MC v C1* | *MC v C3* | *C1 v C3* | *MC v C1* | *MC v C3* | *C1 v C3* | *MC v C1* | *MC v C3* | *C1 v C3* | *MC v C1* | *MC v C3* | *C1 v C3* |
| Step Time | 0.7 | - | - | - | 0.998* | 0.997* | 0.995* | 0.998* | 0.997* | 0.995* | 0.998* | 0.997* | 0.995* |
| Stance Time | 9.8* | 0.002 | 0.001 | 1.000 | 0.991* | 0.992* | 0.991* | 0.991* | 0.992* | 0.991* | 0.987* | 0.988* | 0.991* |
| Swing Time | 12.5* | <0.001 | 0.001 | 1.000 | 0.963* | 0.958* | 0.964* | 0.963* | 0.955* | 0.962* | 0.941* | 0.936* | 0.963* |
| Double Support Time | 9.2* | 0.002 | 0.003 | 1.000 | 0.909* | 0.898* | 0.894* | 0.909* | 0.897* | 0.893* | 0.873* | 0.861* | 0.896* |
| Step Length | 3.4* | 0.102 | 1.000 | 0.202 | 0.949* | 0.973* | 0.932* | 0.947* | 0.970* | 0.932* | 0.941* | 0.971* | 0.927* |
| Gait Speed | 2.4 | - | - | - | 0.975* | 0.986* | 0.966* | 0.974* | 0.985* | 0.966* | 0.972* | 0.986* | 0.964* |

**Figure 3.**
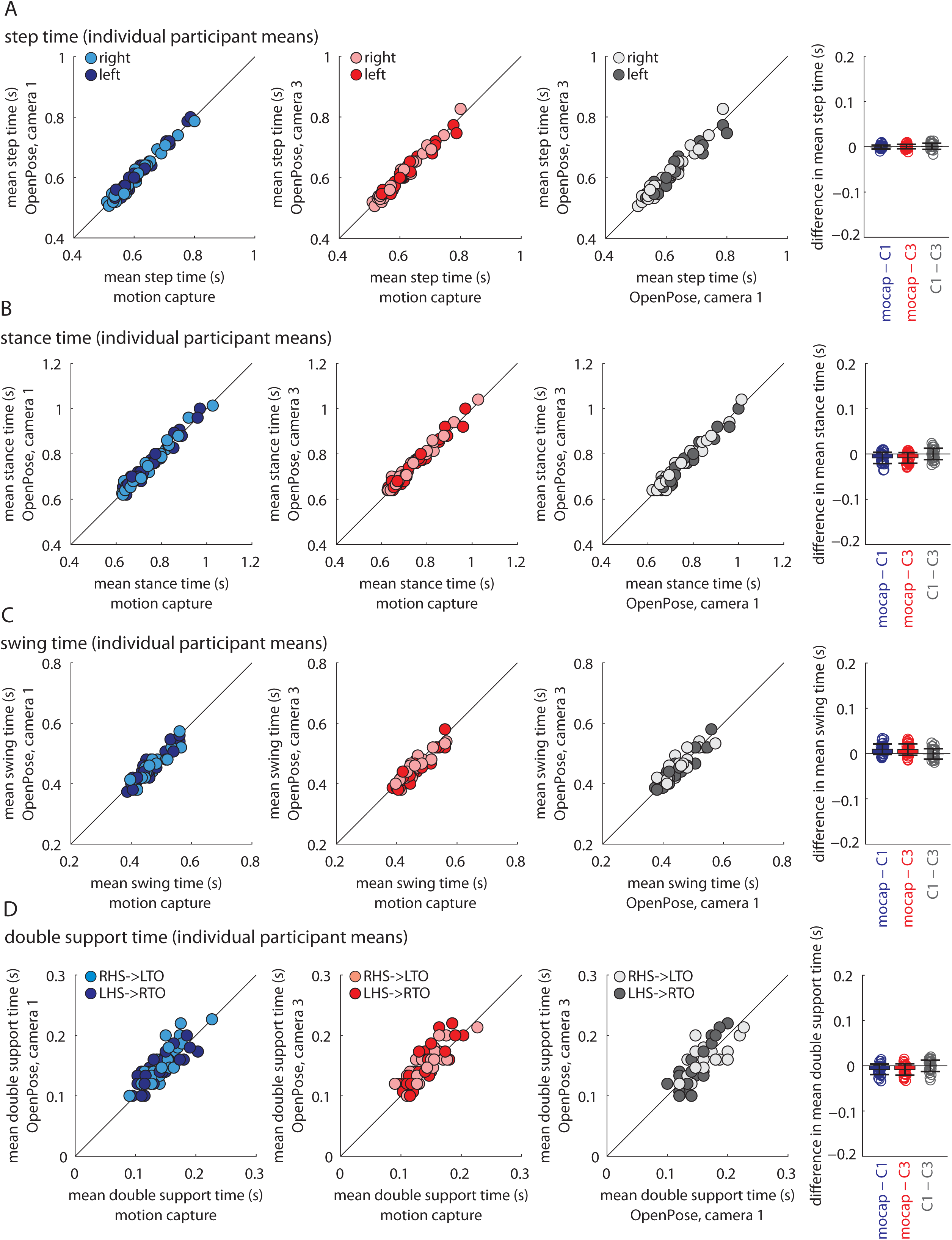
Temporal gait parameters calculated as individual participant means shown for all participants and measurement systems: A) step time, B) stance time, C) swing time, and D) double support time. Bar plots on the far right show individual data, group means, and standard deviations to visualize the distribution of the mean differences observed between the measurement systems. Results from statistical analyses are shown in Table 5.

#### Step length and gait speed

We observed step length and gait speed estimates to be largely similar among the measurement systems. There were no significant differences in step length among the measurement systems when analyzing all steps (Figure 4A) or individual participant means (Figure 4B, Tables 2–5, post-hoc *P*≥0.102). Pearson and intra-class correlation coefficients of step length between motion capture and OpenPose (C1 and C3) were strong: between 0.843 and 0.869 for all steps and ≥0.941 for individual participant means. We also did not observe statistically significant differences when step length was compared between OpenPose left (C1) and right (C3) side views (Tables 3 and 5, post-hoc *P*≥0.202). Pearson and intra-class correlation coefficients of step length between OpenPose left (C1) and right (C3) side views were 0.671 for all steps and ≥0.927 for individual participant means. Similarly, there were no significant differences in gait speed estimates among the measurement systems (Figure 4C), and Pearson and intra-class correlation coefficients were all ≥0.964 (Table 5). All correlations were statistically significant (Tables 3 and 5).

**Figure 4.**
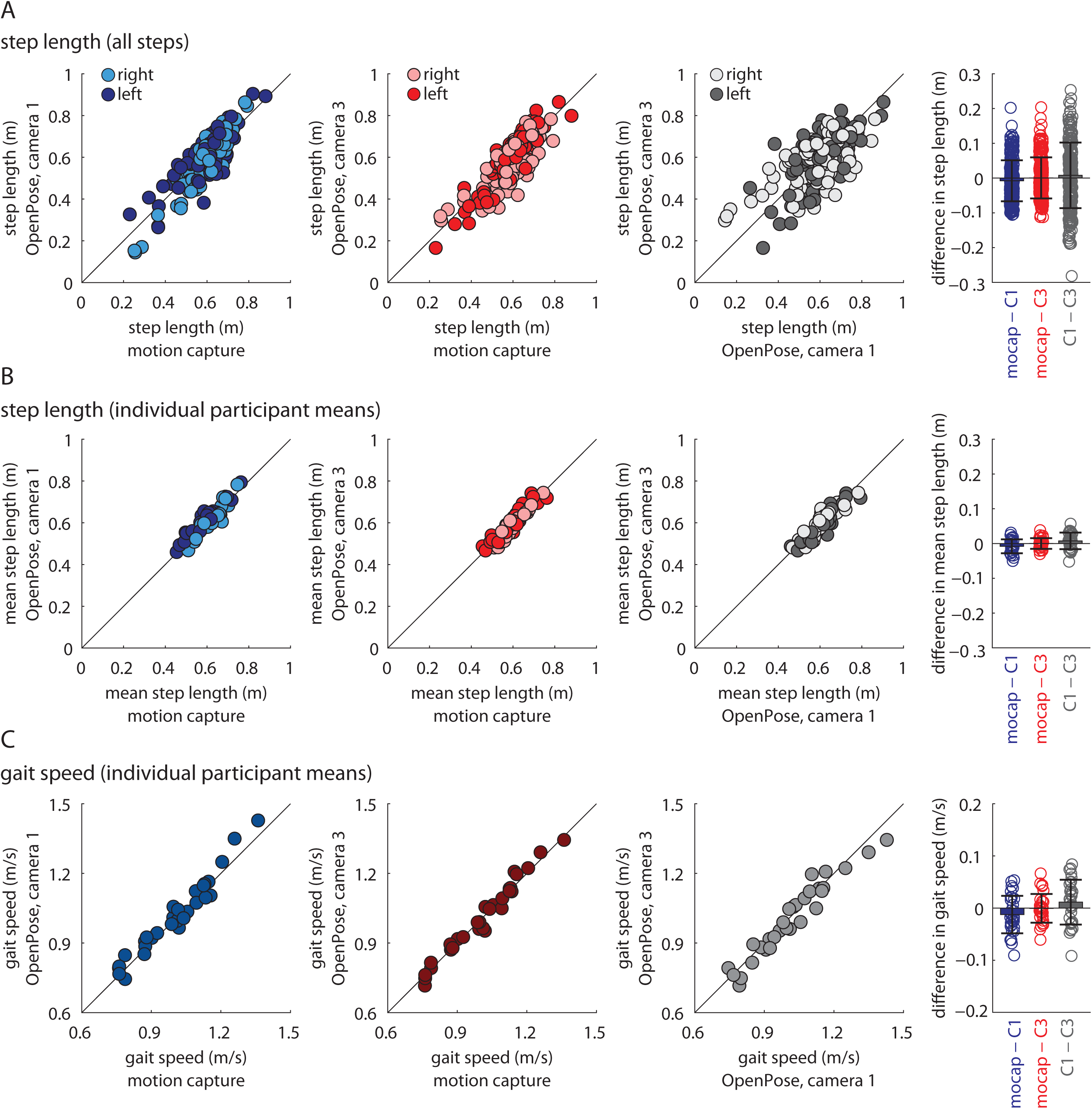
Step length and gait speed comparisons among the different measurement systems. A) Step lengths calculated for all individual steps for all participants and measurement systems. B) Step lengths calculated as individual participant means for all participants and measurement systems. C) Gait speeds calculated as individual participant means for all participants and measurement systems. Bar plots on the far right show individual data, group means, and standard deviations to visualize the distributions of the differences observed between the measurement systems. Results from statistical analyses are shown in Tables 3 and 5.

When examining individual step length data, we occasionally observed large discrepancies between measurement systems. For example, the maximal difference of step length between motion capture and OpenPose (C1 and C3) was 0.204 m, which was substantial given that the length of this step was measured to be 0.672 m by motion capture (i.e., the discrepancy was >30% of the step length). We surmised that anterior-posterior position of the participant on the walkway could have affected how well OpenPose (C1 and C3) estimated step lengths because parallax and changes in perspective could have affected the video-based analyses. We therefore performed a secondary analysis to investigate how anterior-posterior position affected differences in step lengths between the measurement systems.

When step length differences are plotted against anterior-posterior position of the C7 marker (Figure 5), it is apparent that anterior-posterior position affects the step length estimate in OpenPose. When step length is calculated at left heel-strike from a left (C1) side view (Figure 5A, dark circles), step length differences are positive (motion capture greater than OpenPose) at the start of the walkway and differences gradually become negative (motion capture less than OpenPose) as the participant traverses the walkway. Differences in step lengths at right heel-strike between motion capture and OpenPose left (C1) side view show the opposite trend (Figure 5A, light circles): differences are negative (motion capture less than OpenPose) at the start of the walkway and gradually become positive (motion capture greater than OpenPose) at the end of the walkway. Note that differences in step length at left and right heel-strikes overlap and are minimized at the middle of the walkway when the person is in the center of the field of view of the cameras. These effects can be observed in the representative images shown in Figure 6.

**Figure 5.**
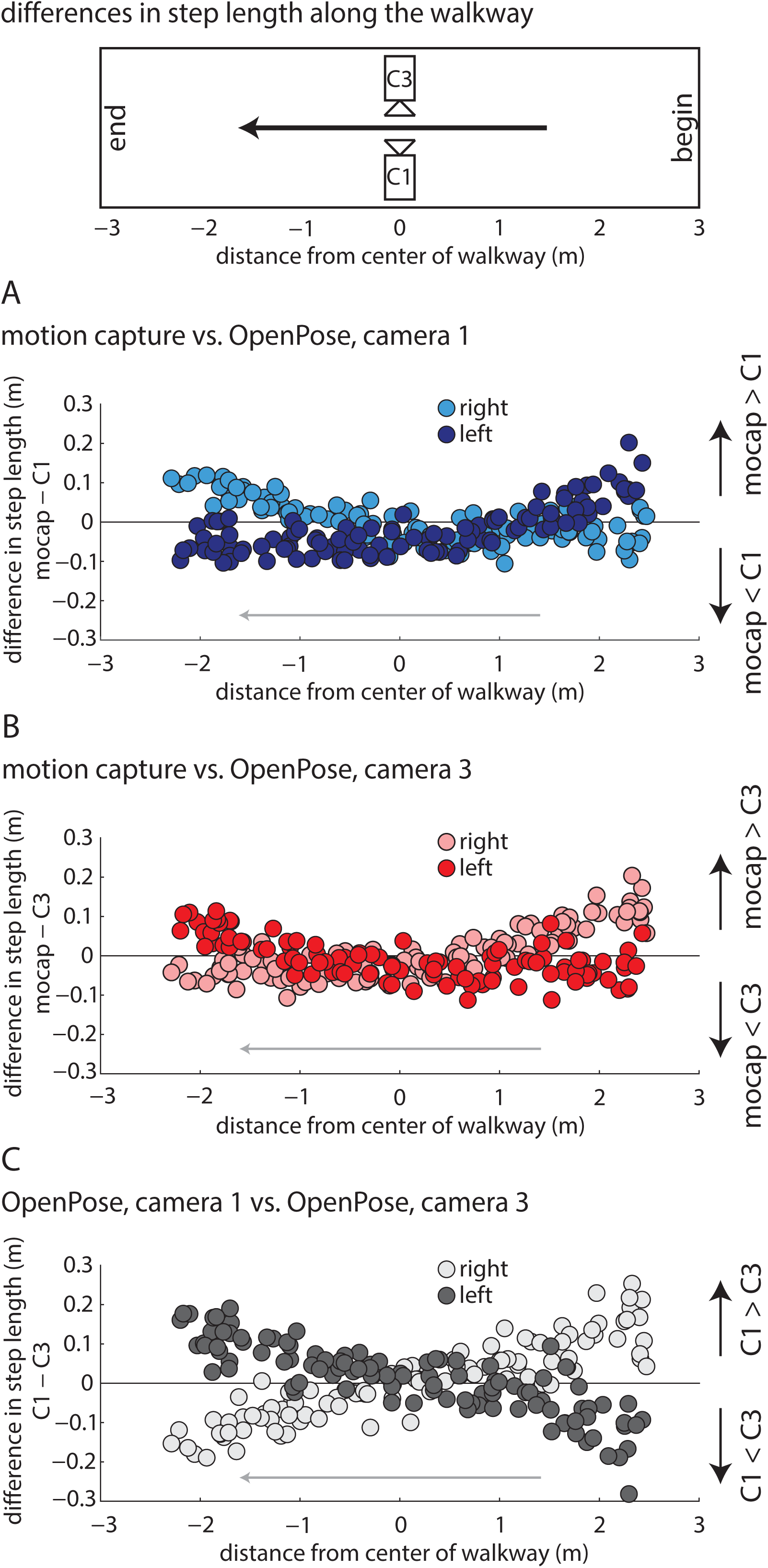
Differences in step lengths calculated from each measurement system in relation to anterior-posterior position on the walkway. Dark circles represent left step lengths, light circles represent right step lengths. A) Differences in step lengths calculated by motion capture and OpenPose left (C1) side views across the walkway. B) Differences in step lengths calculated by motion capture and OpenPose right (C3) side views across the walkway. C) Differences in step lengths calculated by OpenPose left (C1) and right (C3) side views across the walkway.

**Figure 6.**
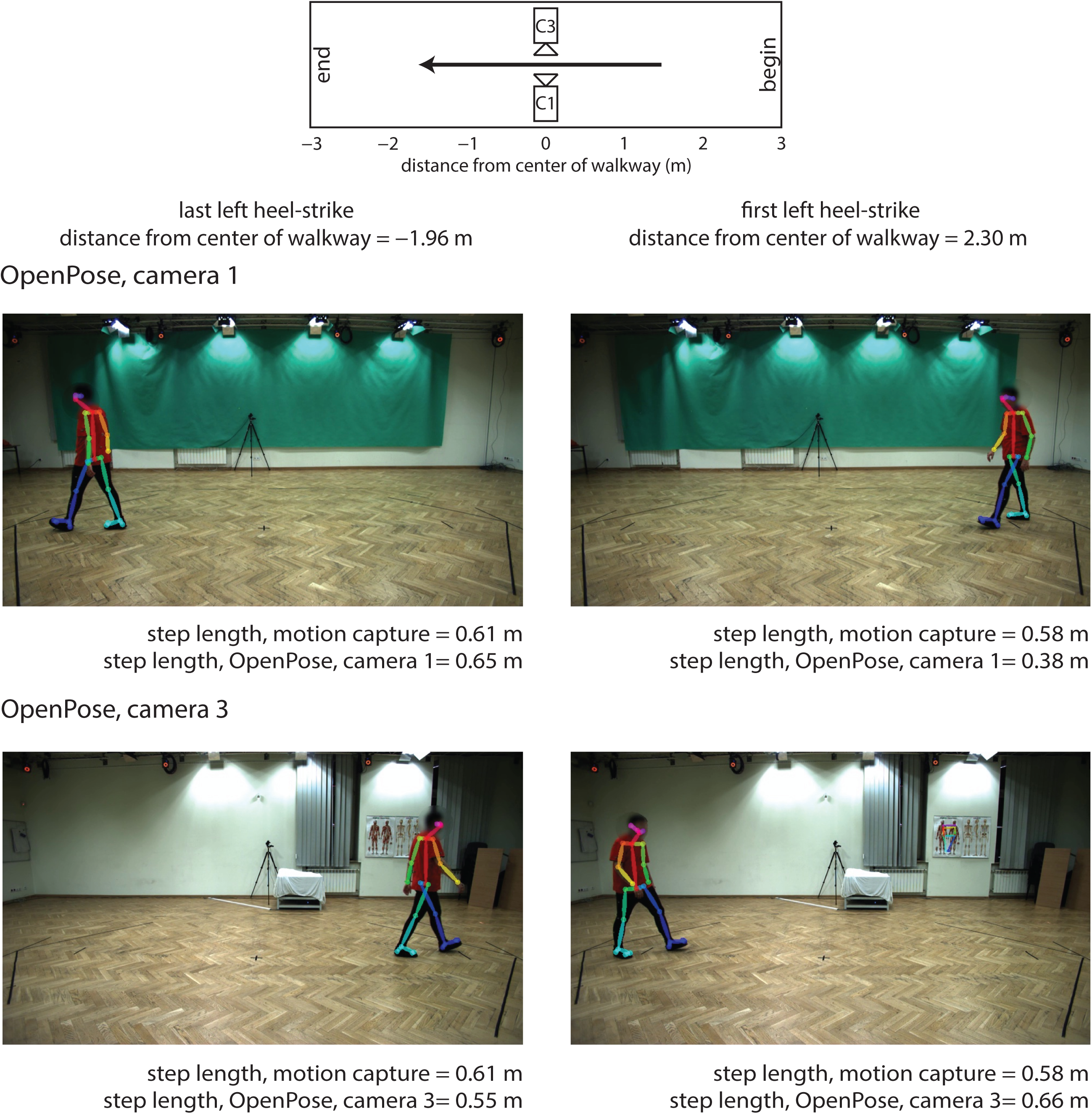
Representative image frames taken from OpenPose output videos highlighting the discrepancies in step length calculation at different positions along the walkway and from different camera views. The public dataset (Kwolek et al, 2019) from which the images belong is made available at http://bytom.pja.edu.pl/projekty/hm-gpjatk/. See Release Agreement for copyrights and permissions.

From the analysis of differences in step length and anterior-posterior position of the participant, we identify the following considerations that affect the estimation of individual step lengths with OpenPose: 1) step length estimation is influenced by the position of the participant along the field of view of the camera, 2) step length estimation is influenced by whether step length is measured at left or right heel-strike, and 3) individual step lengths are estimated most accurately when the person is in the center of the field of view of the camera. If only mean step lengths are of interest for each participant, OpenPose estimates mean step length well because the systematic errors in step length that occur due to position along the walkway appear to offset when step lengths are averaged across the entire walking bout.

### Lower extremity sagittal joint kinematics

Next, we examined how well OpenPose estimated sagittal lower extremity joint angles. We calculated sagittal plane hip, knee and ankle angles across the stride cycle that were averaged for the walking bout of each individual participant (Figure 7). Cross-correlation coefficients (at time lag zero) between motion capture and OpenPose (C1 and C3) were strong for hip and knee angles (all ≥0.975) but less strong (between 0.743 and 0.778) for ankle angles (Table 6). When measuring flexion and extension peaks (or dorsiflexion and plantarflexion peaks at the ankle), we observed a significant main effect of measurement system on peak hip flexion, peak hip extension, peak knee flexion and peak ankle plantarflexion for both legs (Tables 7 and 8). We observed that OpenPose tended to overestimate peak hip flexion and underestimate extension (Figure 8) while differences among systems were mixed in peak knee flexion and peak ankle plantarflexion (Figures 9 and 10). Pearson and intra-class correlation coefficients of peak hip, knee and ankle angles showed large variations between motion capture and OpenPose (C1 and C3) comparisons and between peak angles on left and right legs but were generally not as strong as spatiotemporal parameters: coefficients were between 0.191 and 0.763 for peak hip angles, between 0.113 and 0.541 for peak knee angles and between 0.044 and 0.664 for peak ankle angles (Table 8). Note that a subset of the Pearson and intra-class correlation coefficients did not reach statistical significance (Table 8)

**Table 6.** Cross-correlations of joint angles. *MC*: motion capture; *C1*: OpenPose left side view; *C3*: OpenPose right side view.

|  |  | N | mean±SD |  |  |
| --- | --- | --- | --- | --- | --- |
|  |  |  | *MC* v *C1* | *MC* v *C3* | *C1* v *C3* |
| hip | left | 31 | 0.985±0.013 | 0.976±0.021 | 0.985±0.007 |
|  | right | 31 | 0.975±0.024 | 0.985±0.011 | 0.977±0.018 |
| knee | left | 31 | 0.990±0.006 | 0.986±0.008 | 0.992±0.005 |
|  | right | 31 | 0.982±0.010 | 0.988±0.009 | 0.989±0.008 |
| ankle | left | 31 | 0.774±0.327 | 0.751±0.368 | 0.880±0.119 |
|  | right | 31 | 0.743±0.354 | 0.778±0.303 | 0.898±0.100 |

**Table 7.** Peak joint angles. *MC*: motion capture; *C1*: OpenPose left side view; *C3*: OpenPose right side view.

|  |  |  | N | mean±SD |  |  | mean± SD |  |  | range |  |  |
| --- | --- | --- | --- | --- | --- | --- | --- | --- | --- | --- | --- | --- |
|  |  |  |  | MC | C1 | C3 | MC-C1 | MC-C3 | C1-C3 | MC-C1 | MC-C3 | C1-C3 |
| hip | peak flexion (deg) | left | 31 | 24.1±3.2 | 26.8±2.4 | 26.1±2.5 | -2.7±2.7 | -2.0±3.5 | 0.6±2.0 | [-7.7, 2.8] | [-10.0, 5.1] | [-4.0, 4.2] |
|  |  | right | 31 | 25.0±2.9 | 27.4±3.1 | 27.1±3.1 | -2.3±3.3 | -2.1±3.5 | 0.2±2.9 | [-10.0, 2.1] | [-7.5, 4.7] | [-7.8, 6.3] |
|  | peak extension (deg) | left | 31 | -14.0±3.5 | -16.5±2.6 | -17.5±3.5 | 2.4±2.8 | 3.5±3.8 | 1.1±2.5 | [-3.1, 2.1] | [-3.7, 10.5] | [-3.2, 7.5] |
|  |  | right | 31 | -13.6±3.8 | -18.8±4.2 | -15.8±3.4 | 5.2±3.4 | 2.2±2.5 | -3.0±3.0 | [-0.3, 15.3] | [-1.4, 6.7] | [-10.7, 1.3] |
| knee | peak flexion (deg) | left | 31 | 62.4±5.5 | 64.6±3.7 | 61.5±3.8 | -2.2±4.6 | 0.9±5.2 | 3.1±3.1 | [-10.4, 6.6] | [-8.2, 13.0] | [-2.7, 10.6] |
|  |  | right | 31 | 62.5±5.2 | 60.5±3.2 | 63.6±3.7 | 2.0±4.5 | -1.1±5.1 | -3.1±3.0 | [-7.1, 9.0] | [-13.6, 5.7] | [-8.5, 4.4] |
|  | peak extension (deg) | left | 31 | -1.4±5.2 | 0.0±2.3 | -1.2±2.6 | -1.4±4.5 | -0.2±5.5 | 1.2±2.5 | [-9.3, 7.3] | [-10.4, 11.9] | [-5.6, 5.8] |
|  |  | right | 31 | -1.5±4.7 | -0.8±3.4 | -0.7±3.3 | -0.7±5.1 | -0.8±5.2 | -0.1±2.4 | [-10.4, 7.0] | [-11.3, 8.9] | [-6.9, 3.4] |
| ankle | peak dorsiflexion (deg) | left | 31 | 6.2±7.2 | 6.0±5.1 | 4.3±4.4 | 0.2±7.2 | 2.0±8.3 | 1.7±5.3 | [-13.1, 20.7] | [-10.5, 22.6] | [-11.8, 13.4] |
|  |  | right | 31 | 5.9±8.2 | 5.1±4.3 | 4.7±5.8 | 0.8±8.0 | 1.2±6.2 | 0.4±6.1 | [-12.3, 19.4] | [-11.2, 18.0] | [-11.1, 10.4] |
|  | peak plantarflexion (deg) | left | 31 | -23.7±8.6 | -21.0±4.6 | -27.2±5.4 | -2.6±6.9 | 3.5±9.1 | 6.1±4.2 | [-14.7, 12.2] | [-13.3, 24.8] | [-6.6, 12.6] |
|  |  | right | 31 | -23.4±9.6 | -26.3±7.4 | -22.8±6.0 | 2.9±9.2 | -0.6±7.4 | -3.5±4.9 | [-14.1, 21.2] | [-17.0, 14.4] | [-11.4, 9.2] |

**Table 8.** Peak joint angles. *MC*: motion capture; *C1*: OpenPose left side view; *C3*: OpenPose right side view. Asterisks (*) denote *P* < 0.05.

|  |  |  | *F* | post-hoc (*P*) |  |  | *r* |  |  | ICC<sub>C-1</sub> |  |  | ICC<sub>A-1</sub> |  |  |
| --- | --- | --- | --- | --- | --- | --- | --- | --- | --- | --- | --- | --- | --- | --- | --- |
|  |  |  |  | MC v C1 | MC v C3 | C1 v C3 | MC v C1 | MC v C3 | C1 v C3 | MC v C1 | MC v C3 | C1 v C3 | MC v C1 | MC v C3 | C1 v C3 |
| hip | peak flexion | left | 15.3* | <0.001 | 0.010 | 0.264 | 0.569* | 0.242 | 0.646* | 0.545* | 0.235 | 0.645* | 0.378* | 0.191 | 0.630* |
|  |  | right | 10.0* | 0.001 | 0.006 | 1.000 | 0.402* | 0.325 | 0.569* | 0.401* | 0.325* | 0.569* | 0.310* | 0.265* | 0.575* |
|  | peak extension | left | 20.8* | <0.001 | <0.001 | 0.071 | 0.602* | 0.384* | 0.698* | 0.575* | 0.384* | 0.668* | 0.441* | 0.257 | 0.636* |
|  |  | right | 47.2* | <0.001 | <0.001 | <0.001 | 0.647* | 0.763* | 0.700* | 0.644* | 0.758* | 0.685* | 0.351 | 0.644* | 0.527* |
| knee | peak flexion | left | 8.1* | 0.034 | 1.000 | <0.001 | 0.541* | 0.407* | 0.648* | 0.500* | 0.383* | 0.647* | 0.454* | 0.384* | 0.485* |
|  |  | right | 8.4* | 0.049 | 0.783 | <0.001 | 0.515* | 0.374* | 0.631* | 0.456* | 0.353* | 0.623* | 0.416* | 0.350* | 0.445* |
|  | peak extension | left | 1.9 | - | - | - | 0.473* | 0.139 | 0.505* | 0.355* | 0.113 | 0.501* | 0.341* | 0.117 | 0.453* |
|  |  | right | 0.6 | - | - | - | 0.246 | 0.181 | 0.746* | 0.234 | 0.171 | 0.746* | 0.237 | 0.172 | 0.752* |
| ankle | peak dorsiflexion | left | 1.4 | - | - | - | 0.358* | 0.050 | 0.379* | 0.337* | 0.045 | 0.376* | 0.344* | 0.044 | 0.359* |
|  |  | right | 0.5 | - | - | - | 0.325 | 0.664* | 0.306 | 0.269 | 0.624* | 0.294 | 0.273 | 0.622* | 0.300 |
|  | peak plantarflexion | left | 11.8* | 0.128 | 0.123 | <0.001 | 0.596* | 0.207 | 0.666* | 0.498* | 0.187 | 0.658* | 0.471* | 0.171 | 0.380 |
|  |  | right | 3.9* | 0.284 | 1.000 | 0.001 | 0.437* | 0.637* | 0.747* | 0.423* | 0.572* | 0.731* | 0.408* | 0.578* | 0.651* |

**Figure 7.**
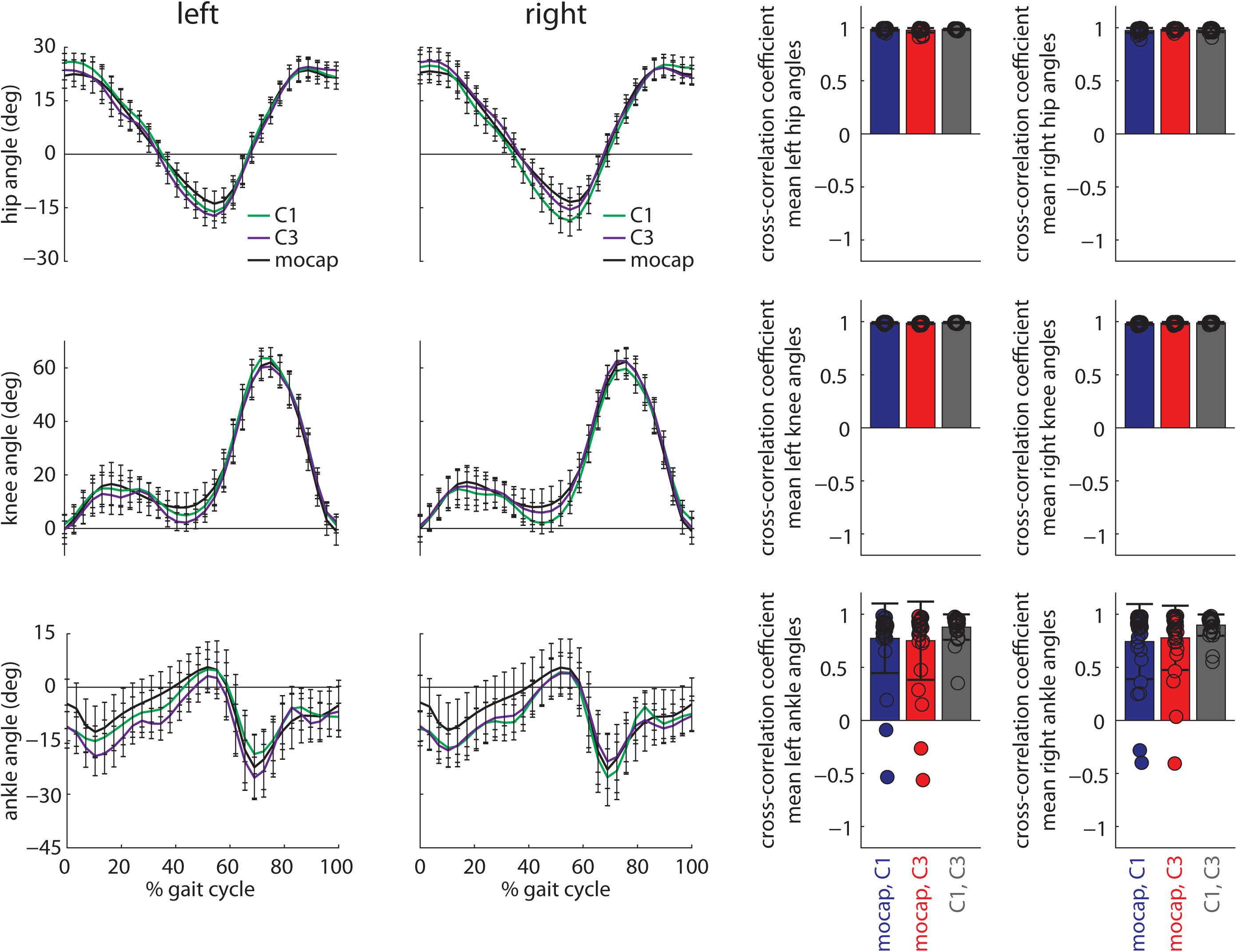
Left) Group mean (±SD) ensemble sagittal joint angles (top: hip, middle: knee, bottom: ankle) for the three measurement systems across both legs. Measurements from motion capture are shown in black, OpenPose left (C1) side view in green, and OpenPose right (C3) side view in purple. For all angles, positive values indicate flexion (or dorsiflexion) and negative values indicate extension (or plantarflexion). Right) Cross-correlation coefficients at time lag zero (individual data, group means, and standard deviations) for individual participant mean joint angle profiles between measurement systems. Color schemes are consistent with Figure 2.

**Figure 8.**
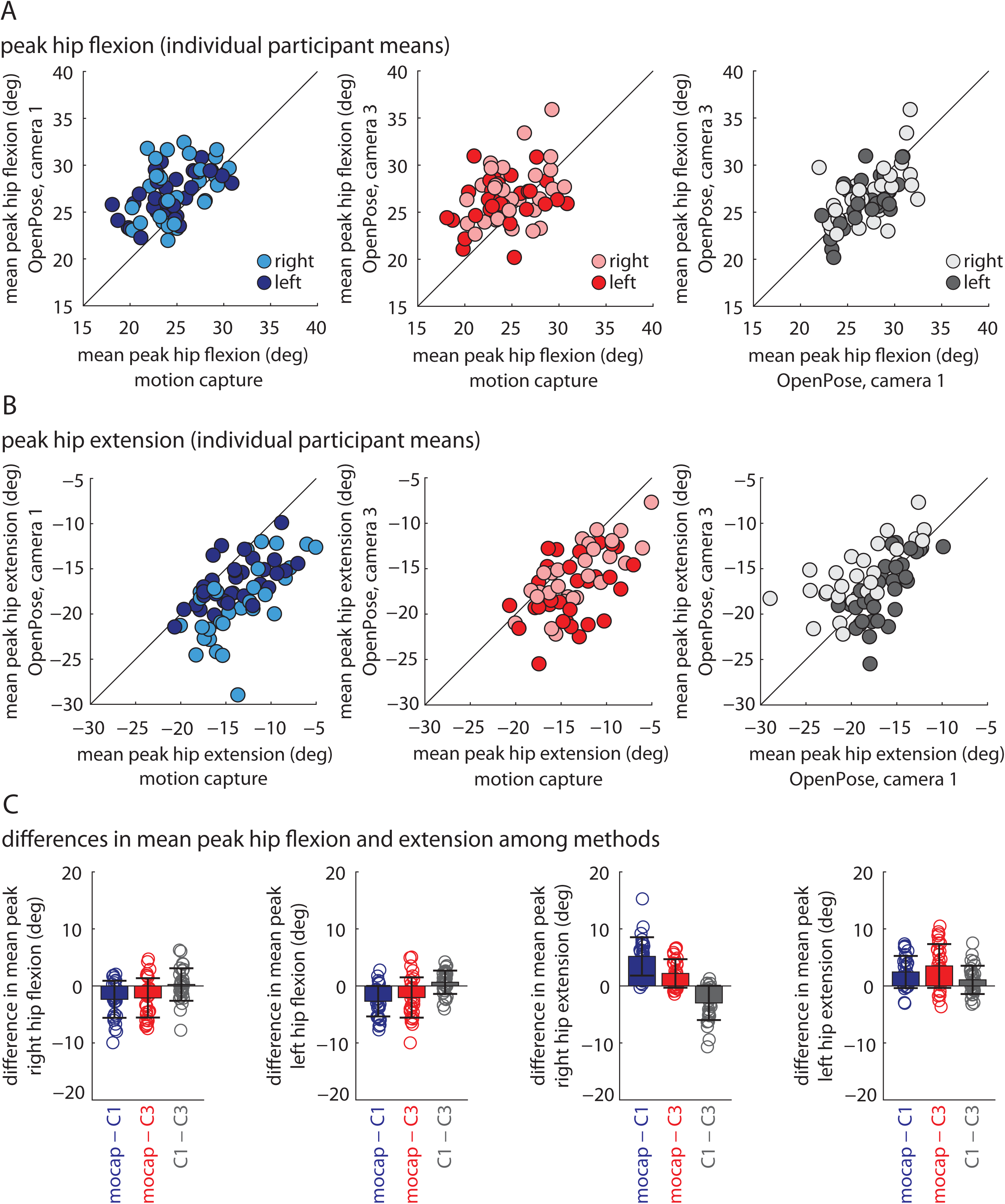
Peak sagittal hip angles calculated as individual participant means shown for all participants and measurement systems: A) peak hip flexion, B) peak hip extension, C) bar plots show individual data, group means, and standard deviations to visualize the distribution of the mean differences observed between the measurement systems. Results from statistical analyses are shown in Table 8.

**Figure 9.**
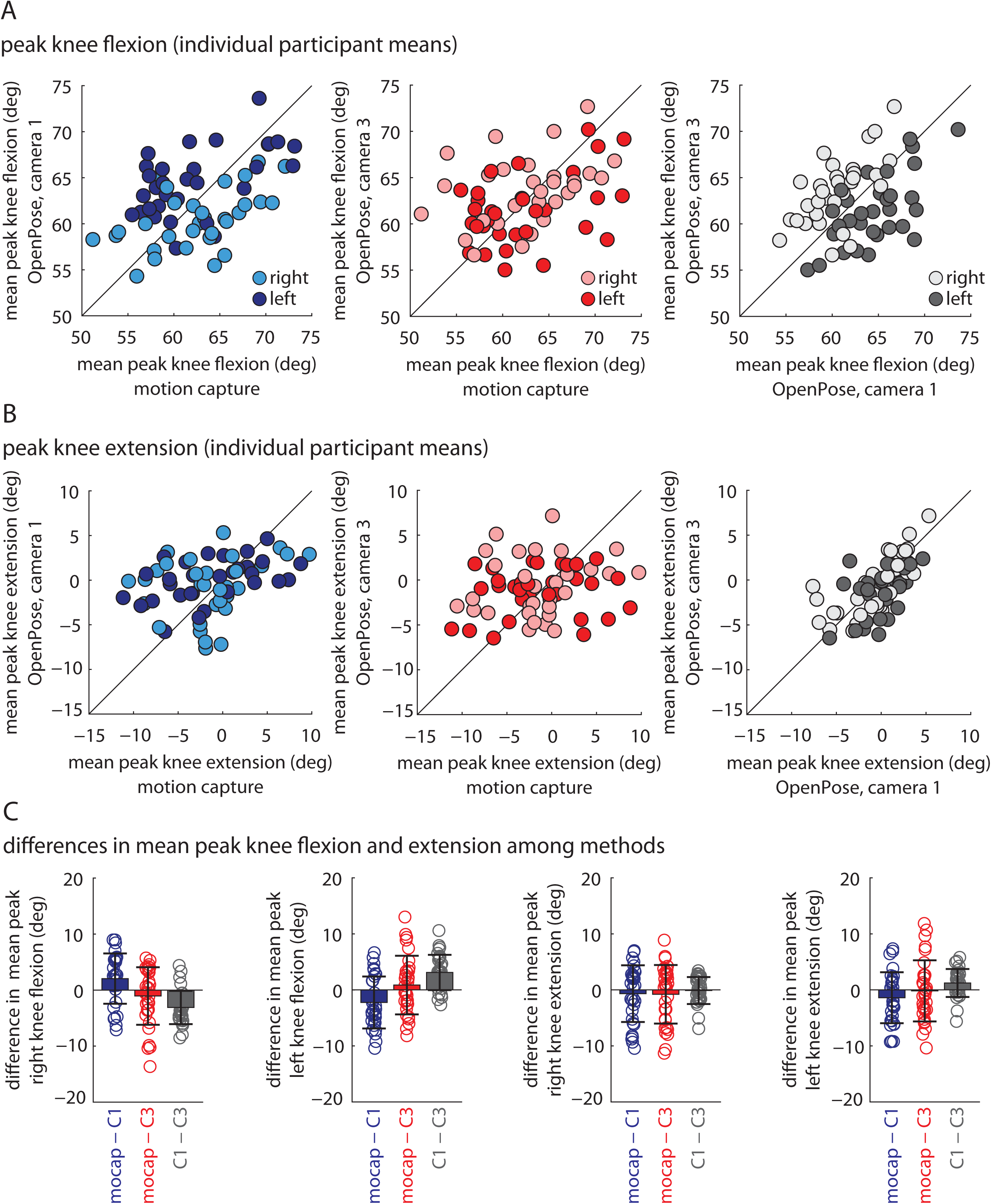
Peak sagittal knee angles calculated as individual participant means shown for all participants and measurement systems: A) peak knee flexion, B) peak knee extension, C) bar plots show individual data, group means, and standard deviations to visualize the distribution of the mean differences observed between the measurement systems. Results from statistical analyses are shown in Table 8.

**Figure 10.**
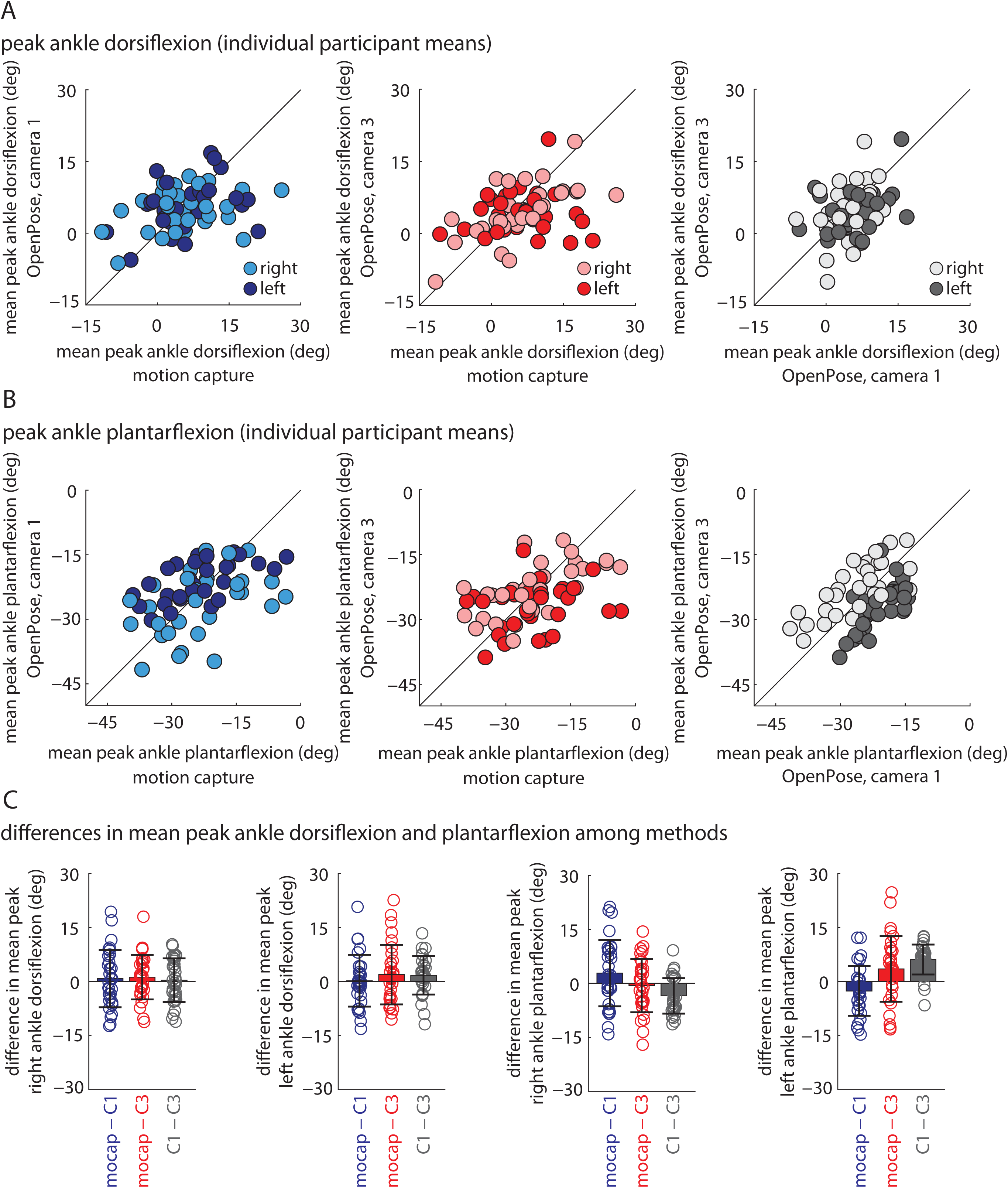
Peak sagittal ankle angles calculated as individual participant means shown for all participants and measurement systems: A) peak ankle dorsiflexion, B) peak ankle plantarflexion, C) bar plots show individual data, group means, and standard deviations to visualize the distribution of the mean differences observed between the measurement systems. Results from statistical analyses are shown in Table 8.

Finally, we compared lower extremity sagittal joint angles between OpenPose left (C1) and right (C3) views. We found relatively strong agreement between the angles calculated from the different views. Cross-correlations between OpenPose left (C1) and right (C3) side views were ≥0.977 for hip and knee angles and between 0.880 and 0.898 for the ankle angle (Figure 7, Table 6). We did observe statistically significant differences between views in peak hip extension, peak knee flexion, and peak ankle plantarflexion (Table 8). Pearson and intra-class correlation coefficients of peak angles between OpenPose left (C1) and right (C3) side views were strongest for the peak hip (between 0.527 and 0.700) and knee angles (between 0.445 and 0.752) and generally weaker for peak ankle angles (between 0.294 and 0.747).

## DISCUSSION

As interest in video-based pose estimation of humans (Andriluka et al., 2014; Toshev and Szegedy, 2014; Insafutdinov et al., 2016; Pishchulin et al., 2016; Martinez et al., 2017; Cao et al., 2019) and animals (Mathis et al., 2018; Nath et al., 2019) increases, there is considerable potential for using these approaches for fast, inexpensive, markerless gait measurement that can be done in the home or clinic with minimal technological requirement. Pose estimation has been used extensively in human gait classification and recognition (Spehr et al., 2012; Kastaniotis et al., 2015; Guo et al., 2017; Kwolek et al., 2019; Sato et al., 2019). Here, we provide initial evidence that pose estimation also shows promise for quantitative spatiotemporal and kinematic analyses of gait that are commonplace in clinical and biomechanical assessments of human walking.

In this study, we aimed to 1) understand how well video-based pose estimation (using OpenPose) could estimate human gait parameters, and 2) provide a workflow for performing human gait analysis with OpenPose. We assessed the accuracy of our OpenPose gait analysis approach by comparing the estimates of spatiotemporal and kinematic gait parameters to measurements obtained by three-dimensional motion capture. We also compared gait parameters estimated by OpenPose analyses from different camera views. We provide a workflow for potential users to determine whether this approach provides data that are satisfactorily accurate for their research or clinical needs and include our interpretations below. Our workflow is provided at https://github.com/janstenum/GaitAnalysis-PoseEstimation.

We observed that OpenPose was able to estimate many gait parameters with a high degree of accuracy. Gait events (e.g., heel-strikes and toe-offs) identified by OpenPose matched those identified by motion capture very well (i.e., generally within one motion capture frame). This led to good performance in the estimation of temporal gait parameters on a step-by-step basis, although motion capture and OpenPose estimations of double support time were less strongly correlated than were the estimations of other temporal parameters (likely due to the relatively short duration of double support periods and 25 Hz frame rate of the cameras).

OpenPose also provided accurate estimates of individual step lengths and participant gait speeds; however, the accuracy of step length estimation worsened when the participant was at the edges of the field of view. Hip and knee kinematics showed good agreement between the motion capture and OpenPose analyses, while OpenPose estimations of ankle kinematics were somewhat less accurate. The accuracy of OpenPose estimations of all gait parameters improved when calculating individual participant mean values (i.e., one summary value of each parameter per participant) rather than discrete step-by-step or peak joint angle values. In sum, our findings show that OpenPose can provide reasonably accurate estimations of many human gait parameters, with some exceptions.

Some prior studies have used OpenPose to investigate particular features of walking or other human movement patterns (Chambers et al., 2019; Sato et al., 2019; Viswakumar et al., 2019; Nakano et al., 2020; Ota et al., 2020; Zago et al., 2020). Our findings align with these reports in that we found OpenPose to be capable of reasonably accurate tracking of human movement (in our case, walking). We also showed that OpenPose estimates of human gait parameters are largely similar across different camera views, an important confirmation since occlusion is often a primary concern when performing two-dimensional movement analysis. We anticipate that in-home and clinical video-based analyses will be performed on videos taken by smartphone, tablets, or other household electronic devices. Many of these devices have standard frame rates of 30 Hz during video recording (and capabilities of up to 240 Hz during slow-motion video recording). The frame rate and resolution of the videos used in our analysis (25 Hz and 960×540, respectively) were lower than the factory settings of most modern smartphones (30 Hz, 1920×1080), suggesting that accuracy may even be improved when using household devices.

Many other markerless motion capture approaches exist for human gait analysis. These include silhouette analyses (Collins et al., 2002; Liang et al., 2003; Lam et al., 2011; Zeng et al., 2014), commercially available products like the Microsoft Kinect (Pfister et al., 2014; Schmitz et al., 2014; Geerse et al., 2015; Kastaniotis et al., 2015; Mentiplay et al., 2015; Xu et al., 2015; Sun et al., 2018), and a variety of other technologies (Corazza et al., 2006; Mündermann et al., 2006; Rhodin et al., 2016; Clark et al., 2019; Kwolek et al., 2019). We did not directly compare the results of our OpenPose analyses to results of any of these other markerless approaches, and thus we are hesitant to speculate about the relative accuracy of our approach against others.

Given the results of several studies cited above, we consider it likely that some of these methods could produce more accurate results. However, a significant advantage of video-based pose estimation with particular relevance for clinic-and home-based gait analysis is that data collection for pose estimation requires no equipment beyond a digital video recording device, whereas many of the other methods require expensive, less accessible, and less portable equipment. We further discuss our observations on the pros and cons of using OpenPose for human gait analysis in the following section.

### Suggestions and limitations

#### Video recordings

Because we did not record the videos used in this study, we did not experiment with different video recording techniques. However, we expect that recording methods with higher frame rates (e.g., the slow-motion video recording feature available on most smartphones), faster shutter speeds, and wider fields of view may improve the accuracy of video-based gait analyses. It is also likely that changes in lighting, clothing, footwear of the participant may affect the ability to track specific keypoints. For example, loose-fitting clothing may introduce more ambiguity into estimations of hip or knee keypoints.

#### Recording in the home or clinic

To generate estimates of gait parameters that incorporate spatial information (e.g., gait speed, step length), it is necessary to scale the video. Here, we accomplished this by scaling the video to known measurements on the ground. This could be done in the home or clinic by placing an object of known size in the field of view at the same depth with which the person is walking.

The videos used in our study were recorded in a large open room with ample distance between the participant and the camera. This allowed recording of several strides per trial and clear sagittal views of the participant. Given the limited availability of large, open spaces in the home or clinic, we perceive a need for approaches capable of estimating at least spatiotemporal gait parameters using frontal plane recordings that are more amenable to narrow spaces.

OpenPose is also capable of three-dimensional human movement analysis. However, this requires multiple simultaneous camera recordings. Because we assumed that most videos taken in the home or clinic will be recorded by a single device, we limited this study to two-dimensional analyses of human walking.

#### Network selection

Here, we used the demo and pre-trained network provided by OpenPose because we considered that this is the most accessible approach that is likely to be used by most new users of OpenPose, especially those who do not have significant expertise in engineering or computer science. Another advantage of using the pre-trained network is that it has already been trained on thousands of images, saving the user significant time and effort in training their own network.

However, it may be possible to obtain more accurate video-based analyses by training gait-specific networks from different views (e.g., sagittal, frontal). Similarly, if a user aims to study clinical populations with gait dysfunction (e.g., stroke, Parkinson’s disease, spinal cord injury) or children, it may be beneficial to train networks that are specific to each population. Furthermore, we did not compare our workflow to other available pose estimation algorithms (e.g., DeeperCut (Insafutdinov et al., 2016), DeepLabCut (Mathis et al., 2018), DeepPose (Toshev and Szegedy, 2014), AlphaPose (Fang et al., 2017)). These approaches are evolving rapidly and are likely to continue to improve in the near future.

#### Potential sources of error in video-based estimates of gait parameters vs. motion capture

We considered several potential reasons for discrepancies between the parameters estimated by OpenPose and those measured using three-dimensional motion capture. Some sources of error may be intrinsic to OpenPose. First and most obvious, OpenPose does not track movements of the human body perfectly from frame-to-frame. Second, the body keypoints identified by OpenPose are unlikely to be equivalent to the marker landmarks. OpenPose relies on visually labeled generalized keypoints (e.g., “ankle”, “knee”) whereas marker placement relies on manual palpation of bony landmarks (e.g., lateral malleolus, femoral epicondyle). Certainly, there may also be some degree of error in the placement of the motion capture markers.

Other sources of error may be introduced by the video recording processes. We observed that parallax and changes in perspective toward the edges of the field of view introduced errors into our OpenPose estimates of parameters that rely on spatial information (e.g., step length).

These errors affected estimates of individual steps but appeared to largely offset one another when estimating mean parameters for each participant. Furthermore, blurring of individual frames due to relatively slow shutter speeds and relatively low frame rates may also contribute to inaccuracies in pose estimation. In the dataset that we analyzed here, participants walked at a relatively slow speed of about 1.0 m s^−1^. Tracking faster walking or running could blur images leading to poorer tracking by pose estimation algorithms.

## CONCLUSIONS

Here, we observed that pose estimation (using OpenPose) can provide reasonably accurate estimates of many human gait parameters when compared to three-dimensional motion capture. The pose estimation approach used in this study requires only a two-dimensional digital video input and outputs a wide array of spatiotemporal and kinematic gait parameters. We identified and discussed features of our approach that we believe may have influenced the accuracy of the OpenPose estimation of particular gait parameters, and we provided a workflow that is available at https://github.com/janstenum/GaitAnalysis-PoseEstimation. We are optimistic about the potential that this initial study reveals for measuring human gait data in the sagittal plane using video-based pose estimation and expect that such methods will continue to improve in the near future.

## ACKNOWLEDGEMENTS

This study was funded by NIH grant R21AG059184 to RTR. We thank Amy Bastian, Darcy Reisman, and their lab members for helpful comments.

